# PhyloBrain atlas: a cortical brain MRI atlas following a phylogenetic approach

**DOI:** 10.1101/2020.07.15.205401

**Authors:** M Zhernovaia, M Dadar, S. Mahmoud, Y Zeighami, Maranzano

## Abstract

Cortical atlases constitute a consistent division of the human cortex into areas that have common structural as well as meaningful and distinctive functional characteristics. The most widely used atlases follow the cytoarchitectonic and myeloarchitectonic characteristics of the cortex and have been combined to the standard anatomical nomenclature of gyri and sulci. More recently, common functional features depicted by resting state functional MRI have also guided the division of the cortical brain in functional regions of interest. However, to date, there are no atlases that divide the cortex considering the common evolutionary changes experienced by the mammalian cortex.

Hence, the present study proposes the division of cortical areas into five main regions of interest (ROIs) following a phylogenetic approach: 1- archicortex, 2- paleocortex, 3- peri-archicortex, 4- proisocortex, 5-neocortex, and twelve neocortical sub-ROIs: 5.1.temporopolar, 5.2.post-central, 5.3.pre-central, 5.4.pericalcarine, 5.5.superior temporal, 5.6.middle temporal, 5.7.precuneus, 5.8.insular, 5.9.inferior parietal, 5.10.caudal anterior, 5.11.posterior cingulate, and 5.12.lingual gyrus.

The segmentations were done using the T1-weighted MNI-ICBM152 non-linear 6th generation symmetric average brain MRI model.

## BACKGROUND AND SUMMARY

Brain atlases aim to divide the organ into a number of regions that have common and meaningful structural and functional characteristics. They constitute a fundamental tool to study and quantify changes in healthy and pathological states. Historically, one of the best known and most used cortical parcellations is the one performed by Brodmann, based on cytoarchitectonic differences of 47 areas, which were considered, at a structural cellular level, the basis of distinct functions (Brodmann and Gary 2006). The Brodmann areas were then translated into modern 3D imaging by Talairach and Tournoux (Talairach, Tournoux et al. 1993) and have been extensively used by neuroscientists investigating pathological conditions linked to changes of specific parcels (Strotzer 2009, Ardila, Bernal et al. 2016, Kawachi 2017, Ueda, Fujimoto et al. 2017). In parallel, cortical myelin architecture has also been studied, generating a cortical parcellation that follows myeloarchitectural differences. The first systematic and best known parcellation of this type was done by the Vogt-Vogt school (Nieuwenhuys 2013). Additionally, cortical parcellation has also been done following a pure gyral anatomical approach, using the standard anatomical nomenclature (Destrieux, Fischl et al. 2010), referring to the corresponding cytoarchitectonic area(s) in a given gyrus, when necessary (e.g. Brodmann area 4 in the pre-central gyrus). Presently, modern histology and magnetic resonance imaging (MRI) have unveiled the limitations of the Brodmann and Vogt parcellations (Amunts and Zilles 2015), and different groups have developed a variety of parcellation systems applying different methods. Over the last two decades, some of the most popular approaches have included: 1) inflation and flattening of the cortex to improve visualization and facilitate the work on this strongly folded structure (Fischl, Sereno et al. 1999) in sulcal and gyral cortices, following the standard anatomical nomenclature (Destrieux, Fischl et al. 2010), 2) inclusion of a wide range of ages and neurodegenerative cases to allow parcellation of cortices into gyri affected by various degrees of atrophy (Desikan, Ségonne et al. 2006), (Klein and Tourville 2012), 3) organization of networks of functionally coupled regions using resting-state functional MRI (Yeo, Krienen et al. 2011), and 4) single subject high-resolution histology, using Nissl stain and immunohistochemistry, correlated with 7TMRI (Ding, Royall et al. 2016) following Brodmann’s and an anatomical gyral parcellation.

To date, there has been no atlas following a phylogenetic approach, a method which would allow the study of the human cerebral cortex according to the developmental events taking place during the millennia of primate evolution. A phylogenetic approach could unveil structural features of different cortical areas grouped on the basis of common evolutionary development.

During mammalian evolution, the archi- and paleocortex, also grouped under the name of allocortex (allocortex=cortex with a number of layers different than six), precedes the development of the larger neocortex. The archicortex is represented by the hippocampus, formed by the Cornus Ammonis areas, dentate gyrus and subiculum, presubiculum, parasubiculum, entorhinal cortex, retrosplenial cortex, and a cortical band in the cingulate gyrus (Mai and Paxinos 2012). The paleocortex, which shares common characteristics with the three-layered cortex of reptiles (Klingler 2017), is composed of the prepiriform and piriform regions, as well as part of the amygdala, olfactory cortex, olfactory bulb, retrobulbar, olfactory tubercle and septal regions (Mai and Paxinos 2012).

The much younger human neocortex (or isocortex) is characterised by six cellular layers depicted by classical Nissl stains. It comprises the sensory areas (somatosensory, auditory, and visual areas), multimodal association areas, and motor areas (Mai and Paxinos 2012).

Finally, the areas of cortices between archi- paleo- and neocortex show a gradual transition of cytoarchitecture, which allow their classification as periarchicortex, adjacent to the archicortex, and proisocortex, between the periarchicortex and neocortex (Fleischhauer 1976) see Figure 1. Our study aimed to construct each phylogenetic cortical parcel by grouping pre-existing areas of known phylogenetic origin on a standard MRI neuroanatomical template in pseudo-Talairach space, useful for morphometry studies (Manera, Dadar et al. 2019).

**Figure 1:**
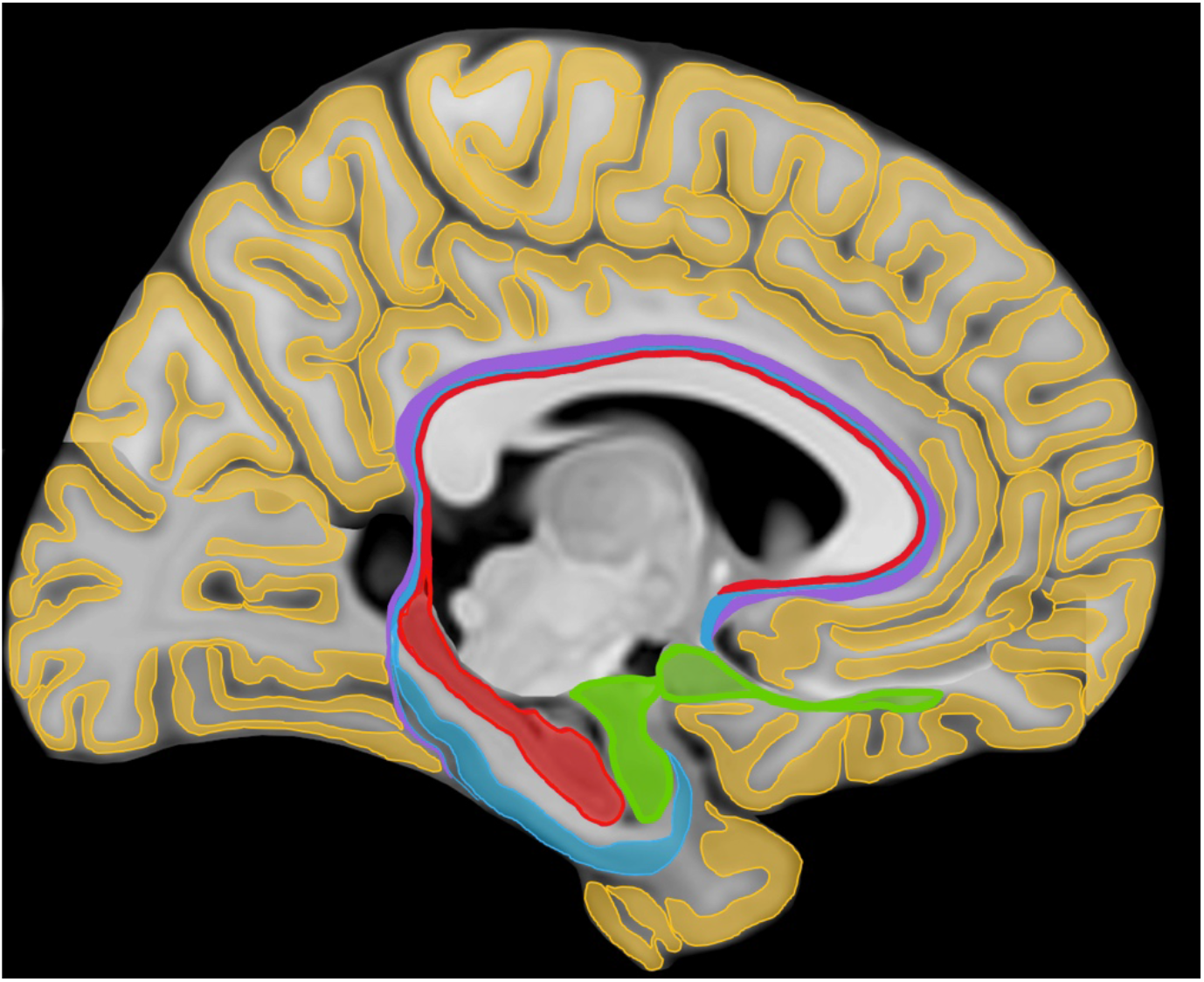
Schematic representation of the subdivisions of the human cerebral cortex into paleocortex (in green), archicortex (in red), periarchicortex (in blue), proisocortex (in purple) and neocortex (in yellow). Modified from Zilles and Amunts, 2012. *Note: we created a modified sagittal representation of the MRI template by fusing various parasagittal slices that allowed the delineation of the five ROIs in the same figure*.

We present the PhyloBrain atlas, which divides the human cerebral cortex into five main regions of interest (ROIs): 1-archicortex, 2-paleocortex, 3-peri-archicortex 4- pro-isocortex, 5-neocortex, and twelve neocortical sub-regions of interest (sub-ROIs), which include various sensory modalities, motor and association cortices: 5.1.temporopolar, 5.2.post-central, 5.3.pre-central, 5.4.pericalcarine, 5.5.superior temporal, 5.6.middle temporal, 5.7.precuneus, 5.8.insular, 5.9.inferior parietal, 5.10.caudal anterior cingulate, 5.11.posterior cingulate, and 5.12.lingual.

## METHODS

The atlas was done following previously described MRI protocols (Pruessner, Li et al. 2000); (Ulmer and Jansen 2013); (Gonçalves Pereira, Insausti et al. 2005); (Frankó, Insausti et al. 2014); (Mikhael, Hoogendoorn et al. 2018); (McCormick, Ziebell et al. 2006); (Palomero-Gallagher, Mohlberg et al. 2008); (Pruessner, Köhler et al. 2002); (Insausti, Juottonen et al. 1998); (Ding, Royall et al. 2016) that described the segmentation of the parcels included within the archicortex (e.g. hippocampi), paleocortex (e.g. piriform cortex (PirC)), peri-archicortex (e.g. perirhinal cortex), proisocortex (e.g. cingulate), and temporopolar neocortex. We performed these segmentations manually on the International Consortium for Brain Mapping of the Montreal Neurological Institute (MNI-ICBM) 2009c average template, which is the most recent version of the MNI-ICBM152 brain average (Manera, Dadar et al. 2019) and provides a higher level of anatomical details. The MNI-ICBM152 non-linear model has two main advantages: 1) it was created from a large number of subjects; hence it represents the average anatomy of the population and is unbiased, unlike single-subject models (Ding, Royall et al. 2016), and 2) the left-right symmetric version enables interpretation of asymmetries that might be found in an analysis.

The sub-ROIs of the neocortical parcels (other than the temporopolar region, which was done fully manual) were based on the manual correction of the CerebrA atlas (Manera, Dadar et al. 2019), by removing voxels of partial volume with the subarachnoideal space cerebrospinal fluid (CSF) and juxtacortical white matter. We chose CerebrA because it was also created on the MNI-ICBM152 2009c average template, providing the perfect complement to our manually created labels.

Both, fully manual segmentations and correction of the CerebrA masks were done using the interactive software package ‘Display’, part of the MINC Tool Kit (https://github.com/BIC-MNI) developed at the McConnell Brain Imaging Center of the Montreal Neurological Institute. This program allows simultaneous viewing and segmentation in the coronal, sagittal and axial planes. Each window allows zooming in and out and a painting tool allows marking voxels with a given color (label/mask number) (Maranzano, Rudko et al. 2016). All the labels created for the PhyloBrain atlas consist of cortical gray matter only.

### ROI 1. Archicortex manual segmentation

we included the hippocampus and entorhinal cortices. We did not attempt the segmentation of the pre- and supra-commissural (indusium griseum) hippocampal portions, because the resolution of the MRI brain template precluded its conclusive segmentation.

The hippocampus is traditionally divided into a posterior portion called the hippocampal tail (HT), a more anterior part called the hippocampal body (HB) and the most anterior, the hippocampal head (HH) (Ulmer and Jansen 2013).

The hippocampus included the dentate gyrus, the cornu ammonis (CA) regions, the part of the fasciolar gyrus that is adjacent to the CA regions. The white matter portions traditionally included in the MRI segmentation of the hippocampus (Pruessner, Li et al. 2000) were excluded, because our atlas aims to exclusively include cortical gray matter. Hence, the fimbria, located at the superomedial level of the HB and the alveus, which separates the HH from the amygdala at the supero-rostral level, were not included in our atlas.

The uncal recess of the inferior horn of the lateral ventricle (LV) and alveus served as landmarks for definition of the supero-anterior border of hippocampus (Frankó, Insausti et al. 2014). The most posterior part of the HT was selected in the coronal plane, where an ovoid mass of gray matter (GM) is first visible inferio-medial to the trigone of the LV (Pruessner, Li et al. 2000); (Ding, Royall et al. 2016). The lateral border of HB was identified by the inferior horn of the LV or the caudally adjacent WM: see Figure 2.

**Figure 2:**
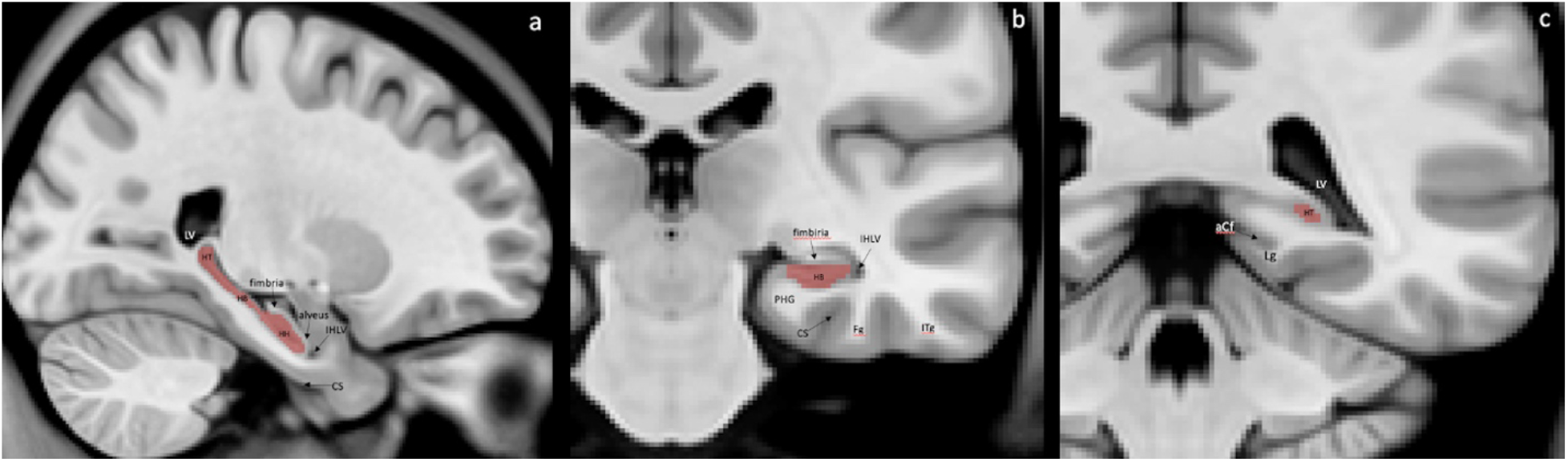
MRI anatomical landmarks of the hippocampal limits. (a) Sagittal section with visualization of anterior, posterior, superior and inferior borders of the hippocampus; delineation of the fimbria that is excluded from the segmentation. (b) Coronal sections of the medial temporal lobe (MTL) with posterior end of the hippocampal body and (c) posterior end of the hippocampal tail. IHLV= inferior horn of lateral ventriculus; LV= lateral ventriculus; CS= collateral sulcus; PHG= parahippocampal gyrus; Fg= fusiform gyrus; Itg= inferior temporal gyrus; aCf= anterior calcarine fissure; Lg= lingual gyrus; HB= hippocampal body; HT= hippocampal tail.

The entorhinal cortex (EC) was segmented by selecting the coronal view and moving from anterior to posterior, from the posterior limit of the temporopolar cortex (described in section ROI.5.1.) towards the point of transition between the HH and HB (intralimbicus gyrus) (Ulmer and Jansen 2013). The collateral sulcus (CS) was localised prior to the segmentation and served as a guide for the subsequent localisation of the first and last slices to be labelled (Huntgeburth and Petrides 2012). If the CS was spreading rostral to the limen insulae (Li), then the anterior border of the EC was taken 2 mm posterior to the first appearance of the Li (grey matter); if the CS was shorter than the Li, then the anterior border was the rostral end of the CS (Pruessner, Köhler et al. 2002); (Insausti, Juottonen et al. 1998). The posterior limit was determined 1 mm posterior to the last slice containing the apex of the intralimbic gyrus (Frankó, Insausti et al. 2014); (Ulmer and Jansen 2013). The superio-medial limit of the EC was given by the sulcus semiannularis (ssa), and the lateral limit was always the midpoint of the medial bank of the CS: see Figure 3.

**Figure 3:**
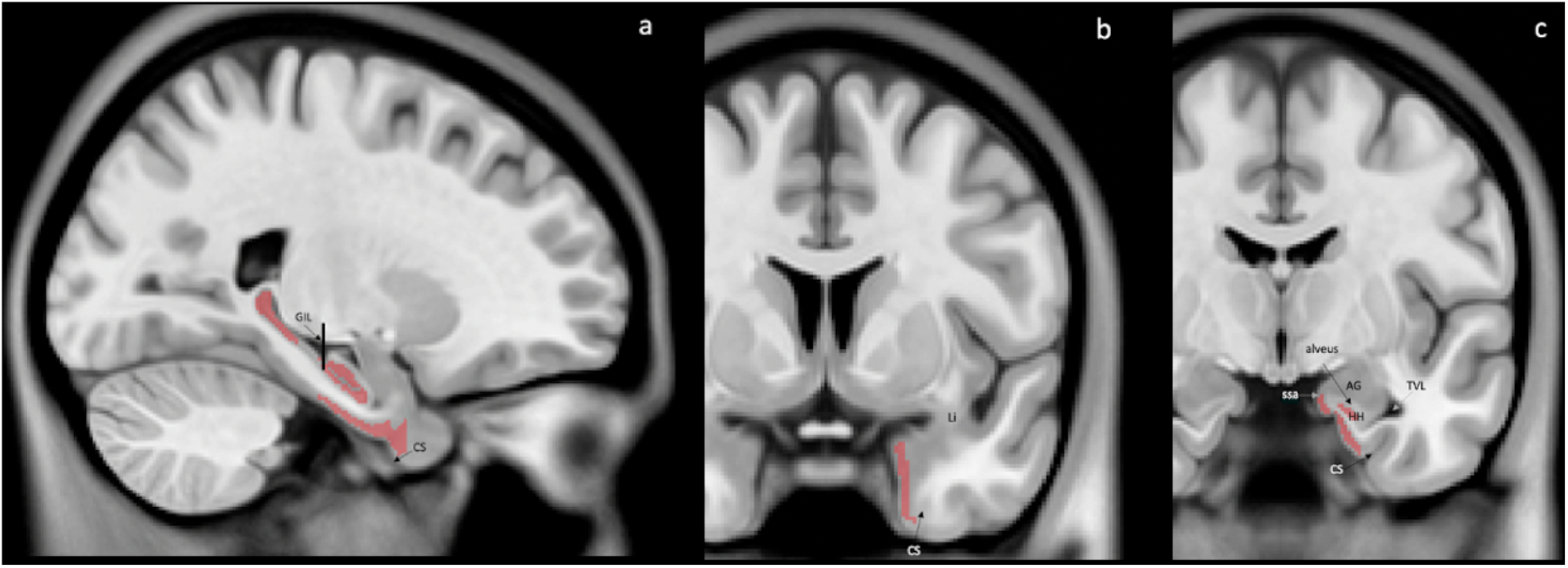
MRI anatomical landmarks of the entorhinal cortex limits. (a) Sagittal section with visualization of anterior and posterior borders of EC. (b) Coronal view of the MTL at the level of the limen insulae and (c) at the level of uncal notch is visible. The perpendicular line indicates the apex of the gyrus intralimbicus. Li= limen insulae GIL= gyrus intralimbicus; CS= collateral sulcus; AG= amygdala; ssa= sulcus semiannularis; TVL= temporal horn of lateral ventricle.

### ROI 2. Paleocortex manual segmentation

The resolution of our MRI precluded the identification of the olfactory bulb and the distinction of the periamygdaloid cortex from the amygdaloid nuclei. Having a close relationship, both anatomically and functionally, the PirC and periamygdaloid cortex together with all the gray matter of the amygdala (AG) were considered in our atlas as a complex (Gonçalves Pereira, Insausti et al. 2005).

The most rostral portion was marked at the level of the Li (white matter-Li in the coronal view) (Pruessner, Li et al. 2000); (Frankó, Insausti et al. 2014).

Superomedially, the fundus of the sulcus semiannularis separates PirC from EC and superolateraly, the fundus of the entorhinal sulcus (es) separates the paleocortex from the substantia innominata. From the level of the Li posteriorly to the rostral end of the AG, the mediolateral extent of the piriform paleocortex occupies progressively more of the surface, (between 30% and 80% of that distance) (Gonçalves Pereira, Insausti et al. 2005), see Figure 4.

**Figure 4.**
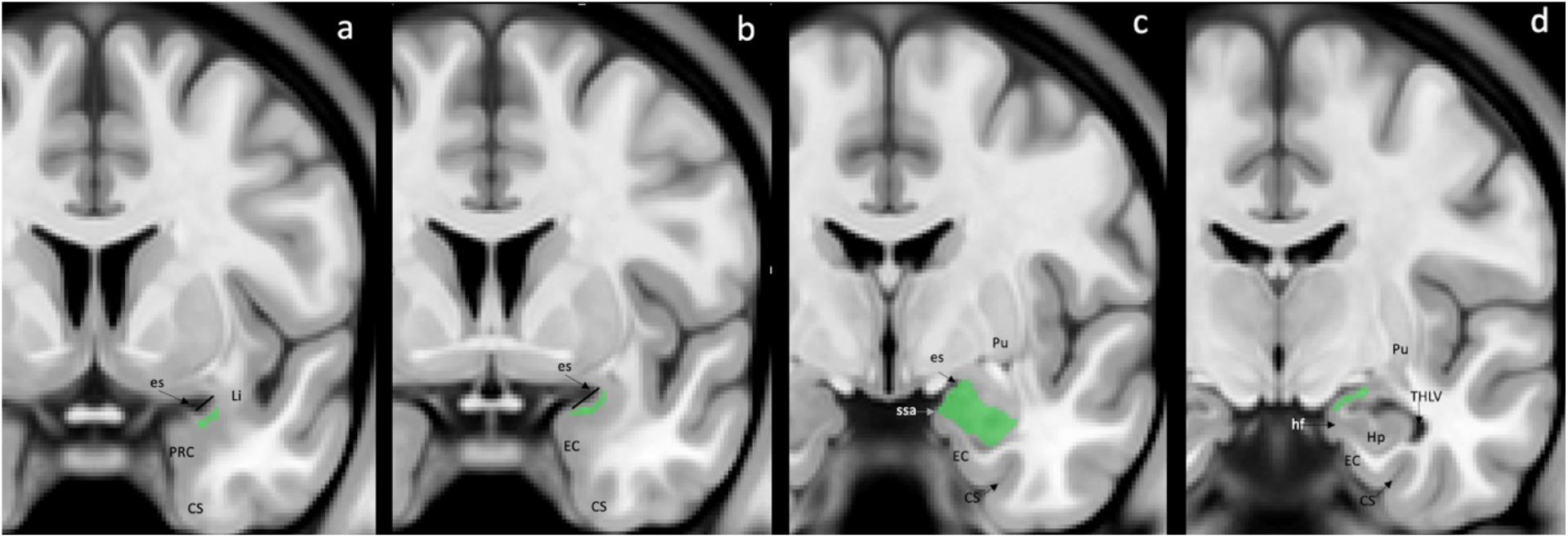
MRI anatomical landmarks of the paleocortex limits. (a) Coronal section of the most rostral part of paleocortex at the level of limen insulae. At this level, piriform-cortical amygdala extends 30% of the distance between the entorhinal sulcus to the most convex point of the medial temporal cortex. (b) The piriform-cortical amygdala occupies 50% of the distance of the entorhinal sulcus – MTL convexity. (c) Coronal section at the level where the piriform-cortical amygdala extends from the entorhinal sulcus down to the sulcus semiannularis. (d) The most caudal section of paleocortex appearance at the level of hippocampal fissure opening. es= entorhinal sulcus; Li= limen insulae; PRC= perirhinal cortex; CS= collateral sulcus; EC= entorhinal cortex; ssa= sulcus semiannularis; Pu= putamen; hf= hippocampal fissure; Hp= hippocampus; THLV= temporal horn of lateral. ventriculus.

Laterally, the gray matter of the AG transitions towards the ventral putamen. The posterior end was defined in the coronal plane at the level of the opening of the hippocampal fissure and at the point where grey matter appears superior to the alveus and laterally to the HH (Pruessner, Li et al. 2000).

### ROI 3. Peri-archicortex manual segmentation

We included the perirhinal and parahippocampal cortices (Ulmer and Jansen 2013). The anterior segment of CS served for determining the rostral limit of PeriAC (Huntgeburth and Petrides 2012). If the CS stretched further anterior than the Li, then anterior tip of the CS was considered as the anterior border of the PeriAC (Frankó, Insausti et al. 2014). If the CS was shorter or as long as the Li, then the border was determined to be 1 mm anterior to it. The most posterior part of parahippocampal cortex was defined as the first posterior slice where the pulvinar was no longer visible (Frankó, Insausti et al. 2014), and it was funnel-shaped, progressively merging with the retrosplenial region. The medial edge stretched from the shoulder of the medial bank of the CS to the medial apex of the parahippocampal gyrus. Laterally, the boundary between the perirhinal and inferotemporal cortices was at the lateral edge of the CS, but if two CS were present, then we considered the fundus of the more lateral CS (Pruessner, Köhler et al. 2002); (Insausti, Juottonen et al. 1998): see Figure 5.

**Figure 5.**
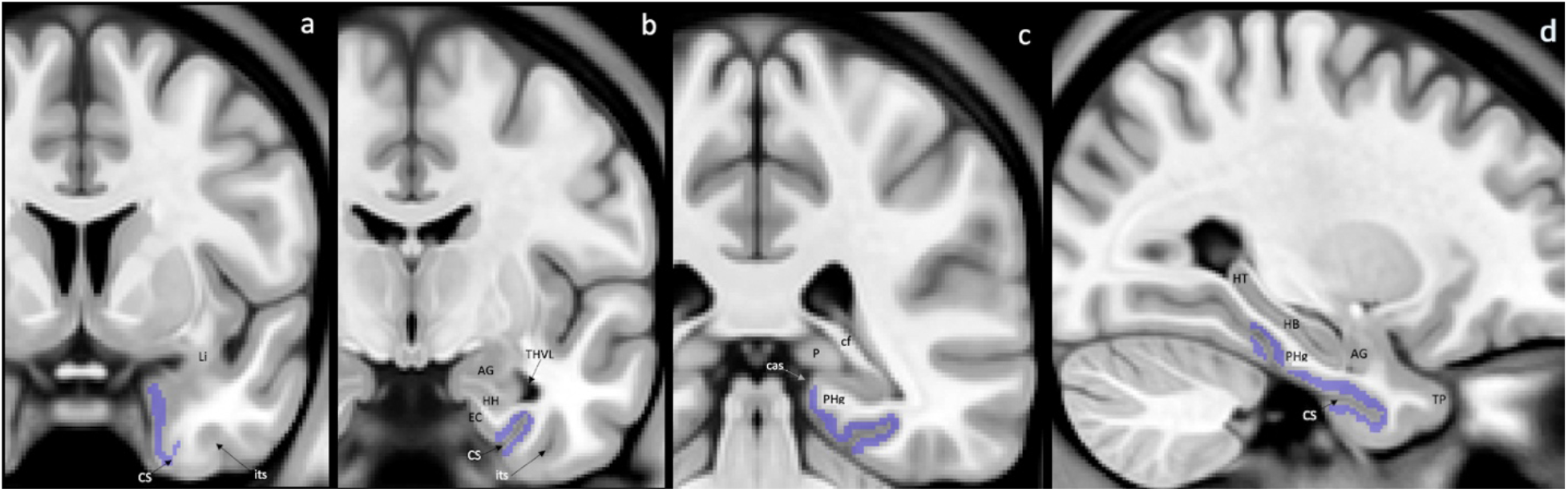
MRI anatomical landmarks of the Peri-archicortex limits. (a) Coronal view of the temporal lobe section where the perirhinal cortex is segmented at the level of the collateral sulcus appearance. (b) Coronal section of the temporal lobe at the level of the perirhinal cortex boundaries with the entorhinal and inferior temporal cortices. (c) Coronal view of the posterior slide where the parahippocampal cortex presents. This is the most posterior level where the pulvinar is present. (d) Sagittal section with visualisation of the peri-archi cortex folded around the collateral sulcus. Li= limen insulae; CS= collateral sulcus; its= inferior temporal sulcus; AG= amygdala; HH= hippocampal head; HB= hippocampal body; HT= hippocampal tail; EC= entorhinal cortex; THVL= temporal horn of ventriculus lateralis; cf= crus of the fornix; P= pulvinar; PHg= parahippocampal gyrus; cas= calcarine sulcus; TP= temporal pole.

### ROI 4. Proisocortex manual segmentation

We included the gray matter of the supra- and sub-callosal areas of the anterior, middle and posterior cingulate gyrus (CinG) (McCormick, Ziebell et al. 2006). In the middle cingulate gyrus, we only considered the proisocortical section (IRd, area infraradiatadorsalis) (Mai and Paxinos 2012) which is immediately adjacent to the location of the indusium griseum and perpendicular to the isocortical part of the cingulate gyrus that occupies the medial surface of the hemisphere (MR, area mediorata) by (Palomero-Gallagher, Mohlberg et al. 2008).

The anterior limit was defined when the CinG was no longer present anterior to the corpus callosum. The gray matter of the CinG around the splenium of the corpus callosum formed the posterior border of proisocortex.

Medially, in the more anterior region, the cortex was limited by gray matter of the superior shoulder of the callosal sulcus (CalS) (gray matter of the CinG), and in the more posterior region by the ventral shoulder of CalS (gray matter of the retrosplenial cortex).

Laterally, the gray matter was segmented up to the deepest point of the bottom of the CalS (for the perigenou part) or at the level of the white matter of the CinG and deepest point of the calcarine fissure for the middle and posterior parts respectively: see Figure 6

**Figure 6.**
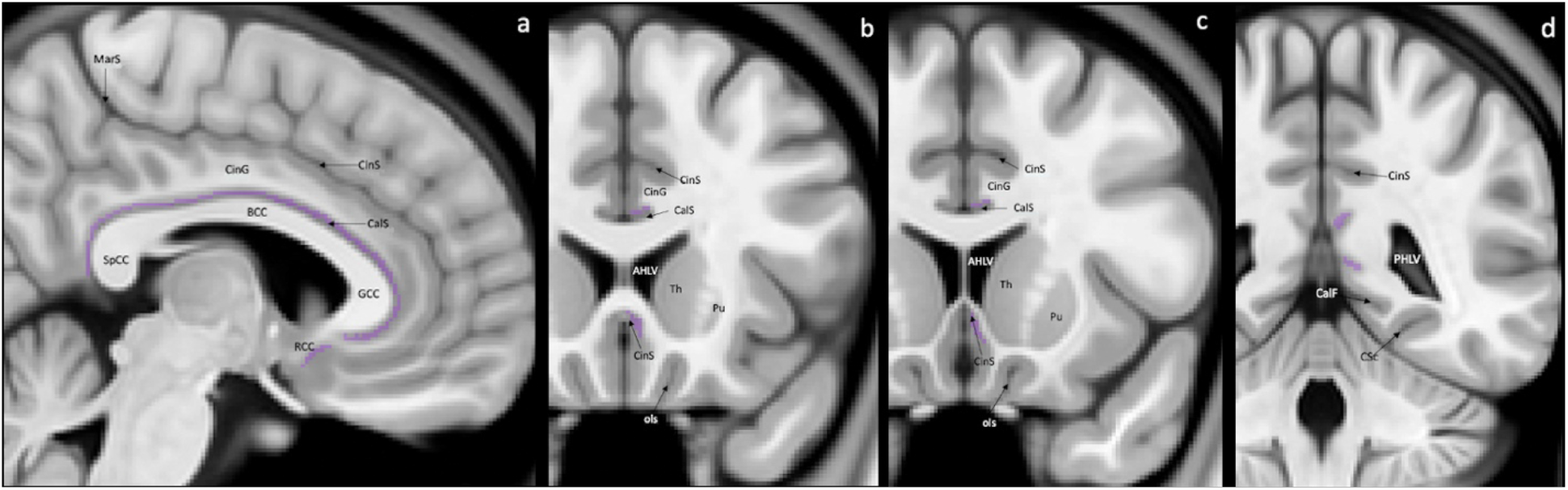
MRI anatomical landmarks of the proisocortex limits. (a) Sagittal view of medial surface of a hemisphere, visualizing supra- and sub-callosal areas of anterior, middle and posterior cingulate gyrus where the proisocortex is present. (b) Coronal section of subcollosal CinC transition to subgenual CinC at the level where the putamen is first visualized within the basal ganglia. (c) Coronal view of the subgenual region at the level of the last slice after which the CinG is no longer present. (d) Coronal section of posterior CinC at the level of calcarine fissure. GCC= genu corpus collosum; BCC= body corpus collosum, SpCC= splenium corpus collosum; RCC= rostrum corpus collosum; CalS= callosal sulcus; CinS= cingulate sulcus; CinG= cingulate gyrus; MarS= marginal sulcus; AHLV= anterior horn of lateral ventriculus; PHLV= posterior horn of lateral ventriculus; Th= thalamus; Pu= putamen; ols= olfactory sulcus; CalF= calcarine fissure; CSc= collateral sulcus caudal segment.

#### ROI. 5.1. Temporopolar neocortex manual segmentation

The anterior limit was determined as the most prominent part of the temporal pole where the grey matter of the temporal cortex is first visible in the middle cranial fossa (Frankó, Insausti et al. 2014). The visualization of the CS or gray matter of the Li defined the posterior border. The superolateral end was determined by the fundus of the temporo-polar sulcus (tps) and thus, the gyrus of Schwailbe (Ulmer and Jansen 2013) and the inferolateral limit was given by the medial bank of inferior temporal sulcus (Pruessner, Köhler et al. 2002); (Insausti, Juottonen et al. 1998): see Figure 7.

**Figure 7.**
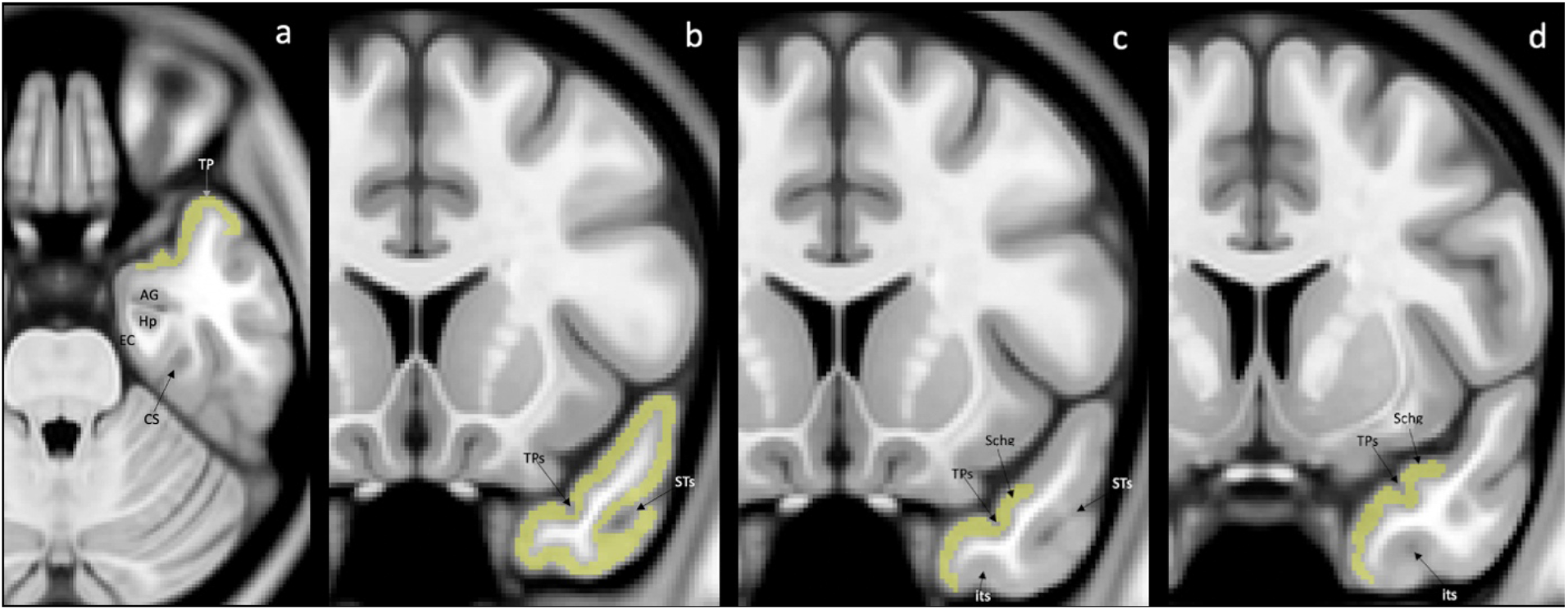
MRI anatomical landmarks of the temporopolar neocortex limits. (a) Axial section of temporal lobe with visualisation of most prominent part of temporal pole. (b)The coronal view of anterior section of temporal lobe, when the inferior temporal sulcus was not yet visible and the section (c) where inferolateral and superolateral borders can be found. (d) The most posterior coronal section of temporopolar neocortex presence before limen insulae gray matter appearance. TP, temporal pole; AG, amygdala; Hp, hippocampus; EC, entorhinal cortex; CS, collateral sulcus; Schg, gyrus of Schwalbe; TPs, temporopolar sulcus; its, inferior temporal sulcus; STs, superior temporal sulcus.

#### ROI 5.2. Post-central gyral cortex (somatosensory)

Corresponding to the somatosensory region, which was modified from the CerabrA atlas (Manera, Dadar et al. 2019), by removing partial volume voxels adjacent to CSF and WM of the original masks labeled as 13 (right hemisphere) and 64 (left hemisphere) of the CerebrA atlas. The editing of the CerebrA mask was done using Display, which allowed saving our final PhyloBrain mask of the post central gyral cortex.

#### ROI 5.3. pre-central cortex (motor)

The original mask of the CerebrA atlas (Manera, Dadar et al. 2019) was uploaded using Display and all the voxels of partial volume with either the adjacent juxtacortical white matter or the sulci cerebrospinal fluid were removed for regions 35 (right hemisphere) and 86 (left hemisphere) from CerebrA. The corrected mask was saved as the final PhyloBrain atlas label for the pre-central gyrus.

#### ROI 5.4. pericalcarine cortex (visual)

A similar procedure as that described in point 5.3. was performed with the original masks labeled 6 (right hemisphere) and 57 (left hemisphere).

#### ROI 5.5. superior temporal cortex (auditory)

A similar procedure as that described in point 5.3. was performed with the original masks 45 (right hemisphere) and 96 (left hemisphere).

#### ROI 5.6. middle temporal cortex

A similar procedure as that described in point 5.3. was performed with the original masks 28 (right hemisphere) and 79 (left hemisphere).

#### ROI 5.7. precuneus cortex (association)

A similar procedure as that described in point 5.3. was performed with the original masks 31 (right hemisphere) and 82 (left hemisphere).

#### ROI 5.8. insular cortex

A similar procedure as that described in point 5.3. was performed with the original masks 23 (right hemisphere) and 74 (left hemisphere).

#### ROI 5.9. inferior parietal cortex

A similar procedure as that described in point 5.3. was performed with the original masks 10 (right hemisphere) and 61 (left hemisphere).

#### ROI 5.10. caudal anterior cingulate cortex

A similar procedure as that described in point 5.3. was performed with the original masks 30 (right hemisphere) and 81 (left hemisphere).

#### ROI 5.11. posterior cingulate cortex

A similar procedure as that described in point 5.3. was performed with the original masks 47 (right hemisphere) and 98 (left hemisphere).

#### ROI 5.12. lingual gyrus cortex

A similar procedure as that described in point 5.3. was performed with the original masks 12 (right hemisphere) and 63 (left hemisphere).

Figure 8 illustrates the different masks identifying the ROIs in a coronal section of the MNI-ICBM152 2009c T1-weighted average template

**Figure 8:**
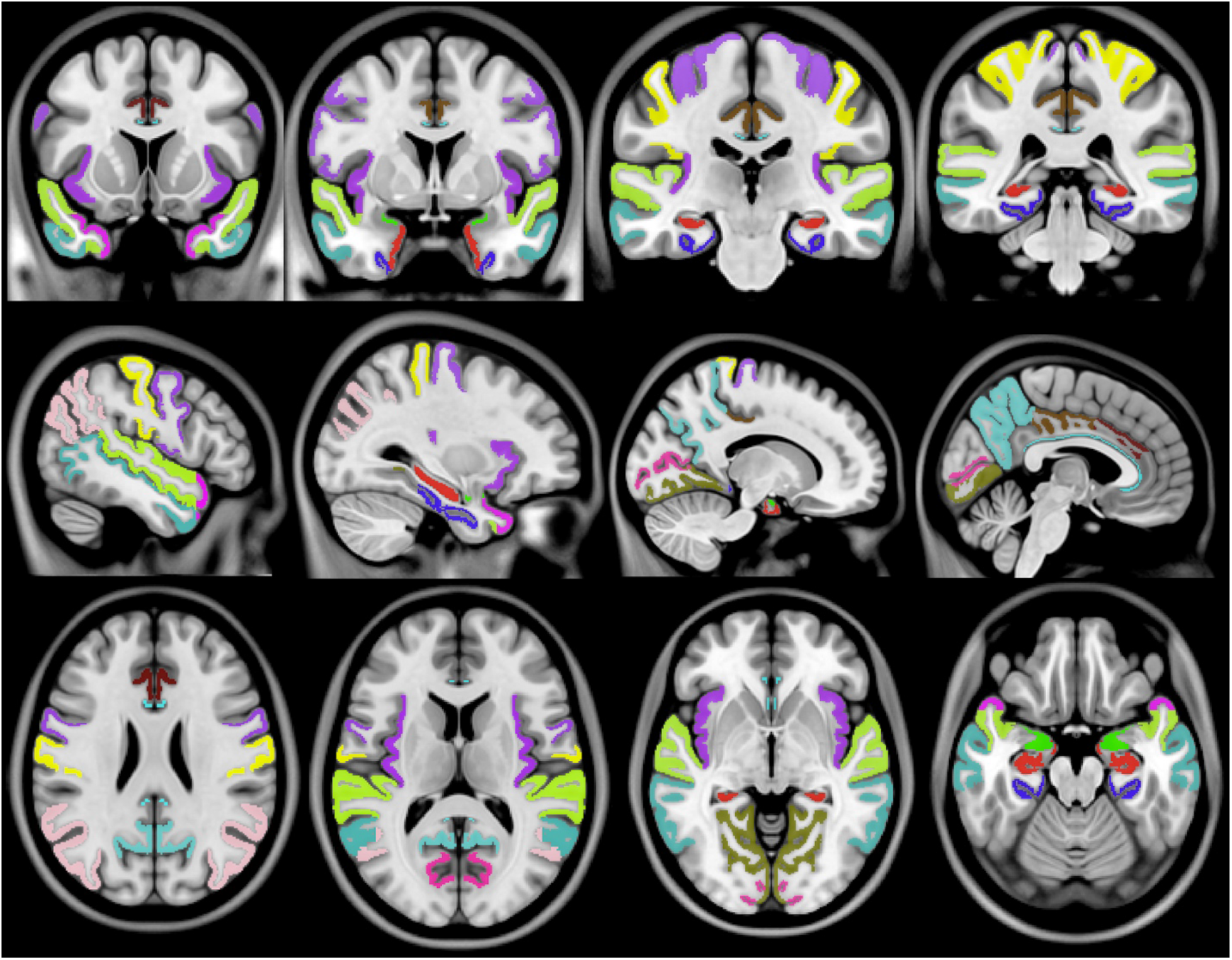
From left to right, various anterior to posterior coronal slices (top row), right to left sagittal slices (middles row), and superior to inferior axial slices (bottom row) of the templates and masks of the ROIs.

## DATA RECORDS AND CODE AVAILABILITY

The final masks of each ROI of the PhyloBrain atlas are available from the corresponding author upon request and also at https://gin.g-node.org/Maryna.Zhernovaia/MRI/. Table 1 indicates the numbers of the MINC format masks of each ROI, in the right and left sides, and the number of voxels (volume in cubic milimeters) of each ROI.

**Table 1:** ROIs names and numbers of the final labels in MINC format

| **LABEL NAME** | **PhyoBrain ID**<br>**Right Hemisphere**<br>**labels** | **PhyoBrain ID**<br>**Left Hemisphere**<br>**labels** | **Number of voxels:**<br>**volume in mm<sup>3</sup>** |
| --- | --- | --- | --- |
| archicortex | 1 | 21 | 3312 |
| paleocortex | 2 | 22 | 1134 |
| periarchicortex | 3 | 23 | 2238 |
| proisocortex | 4 | 24 | 596 |
| temporopolar cortex | 5 | 25 | 3752 |
| post-central cortex | 6 | 26 | 12136 |
| pre-central cortex | 7 | 27 | 11989 |
| pericalcarine cortex | 8 | 28 | 3015 |
| superior temporal cortex | 9 | 29 | 14868 |
| middle temporal cortex | 10 | 30 | 17509 |
| precuneus cortex | 11 | 31 | 12736 |
| insular cortex | 12 | 32 | 7617 |
| inferior parietal cortex | 13 | 33 | 16518 |
| caudal anterior cingulate | 14 | 34 | 1588 |
| posterior cingulate | 15 | 35 | 3016 |
| lingual gyrus cortex | 16 | 36 | 8381 |

The T1w template is available at http://nist.mni.mcgill.ca/?p=904. The imaging data is available in compressed MINC (Vincent, Neelin et al. 2016) and NIfTI formats.

## VALIDATION

### Intra-rater variability assessment

The manual segmentations of the archicortex, paleocortex, periarchicortex, proisocortex, and temporopolar cortex were performed by M.Z. and revised by an expert anatomist (J.M.) with more than 15 years of experience in segmentation of the brain structures on MRI scans. The correction of the CerebrA masks that were used as the starting point for the various neocortical areas considered in this atlas were performed by J.M.

All segmentations (either fully manual or the correction of pre-existing masks) were done twice, to allow the assessment of the intra-rater variability using Dice Kappa similarity index, which determines the proportion of voxels that are common (in the same special location) to the two masks. A Dice Kappa of 1 indicates a perfect spatial overlap, whereas a Dice Kappa of 0 implies no overlap between the two masks.

The Dice Kappa values of the different ROIs were as follows: 0.85 for the archicortex, 0.81 for the paleocortex, 0.69 for the periarchicortex, 0.64 for the proisocortex, 0.80 for the neocortical temporopolar cortex, 0.80 for the neocortical post-central gyrus, 0.93 for the neocortical pre-central gyrus, 0.87 for the neocortical pericalcarine region, 0.98 for the neocortical superior temporal gyrus, 0.94 for the neocortical middle temporal gyrus, 0.79 for the neocortical pre-cuneus region, 0.81 for the insular cortex, and 0.80 for the neocortical inferior parietal gyrus, indicating excellent inter-rater agreement.

## Abbreviations

AG: amygdala
AHLV: anterior horn of lateral ventricle
aCf: anterior calcarine fissure
BCC: body corpus collosum
CA: cornu ammonis
cas: calcarine sulcus
CalS: callosal sulcus
CinG: cingulate gyrus
CinS: cingulate sulcus
cf: crus of the fornix
CS: collateral sulcus
EC: entorhinal cortex
es: entorhinal sulcus
Fg: fusiform gyrus
GIL: gyrus intralimbicus
hf: hippocampal fissure
HB: hippocampal body
HT: hippocampal tail.
HH: hippocampal head
Hp: hippocampus
IHLV: inferior horn of lateral ventriculus
itg: inferior temporal gyrus
its: inferior temporal sulcus
LV: lateral ventriculus
Li: limen insulae
Lg: lingual gyrus
MarS: marginal sulcus
MRI: magnetic resonance imaging
MTL: medial temporal lobe
ols: olfactory sulcus
PHg: parahippocampal gyrus
PeriAC: peri-archicortex
PRC: perirhinal cortex
PirC: piriform cortex
PHLV: posterior horn of lateral ventricle
Pu: putamen
P: pulvinar
RCC: rostrum corpus collosum
ROIs: regions of interest
Schg: gyrus of Schwalbe
SpCC: splenium corpus collosum
ssa: sulcus semiannularis
STs: superior temporal sulcus
Th: thalamus
TP: temporal pole
TPs: temporopolar sulcus
TVL: temporal horn of lateral ventricle
T1w: T1-weighted

## AUTHOR CONTRIBUTIONS

**Maryna Zhernovaia:** Study concept and design, manual segmentation of the labels, analysis and interpretation of the data, drafting and revision of the manuscript.

**Mahsa Dadar**: Study concept and design, analysis and interpretation of the data, revising the manuscript.

**Sawsan Mahmoud**: analysis and interpretation of the data, revising the manuscript.

**Yashar Zeighami**: analysis and interpretation of the data, revising the manuscript.

**Josefina Maranzano**: Study concept and design, manual correction of the labels and supervision of the manual segmentations, analysis and interpretation of the data, drafting and revision of the manuscript

